# Emphasis on quality in iNaturalist plant collections enhances learning and research utility

**DOI:** 10.1101/2023.01.24.525445

**Authors:** Mason C. McNair, Chelsea M. Sexton, Mark Zenoble

**Affiliations:** Clemson University, 2200 Pocket Road, PeeDee Research & Education Center, Florence, SC 29506; University of Georgia, 120 Carlton Street, Athens, GA 30602

## Abstract

Following the switch to remote online teaching in the wake of the COVID-19 pandemic, the plant taxonomy course at the University of Georgia (UGA) switched to iNaturalist for the specimen collection portion of the course requirements. Building off extant rubrics, the instructors designed project guidelines for a fully online plant collection experience to alleviate plant awareness disparity. Researchers collected stratified samples from the UGA iNaturalist project along with four other institutions’ projects to determine if rubrics and project guidelines could improve the quality of observations to make them useful in plant science research. The specific rubric was shown to improve quality of iNaturalist observations. Researchers found that iNaturalist increased engagement as a student-centered tool but did not enhance their manual keying skills as the app uses automatic identification. Instructors recommend continuing to use iNaturalist to supplement physical collection and keying along with a detailed rubric and guidelines for collection.

## Introduction

Modern plant taxonomy utilizes both phenotypic and genotypic data to group populations of plants into understandable groupings which we call species. Teaching students to appreciate this classification process is challenging and was made more difficult during the global COVID-19 pandemic. The Plant Taxonomy course at the University of Georgia (UGA) aims to enable students to explain the importance of plant taxonomy in the context of conservation, systematics, and species concepts. Within this goal, the students will also correctly apply plant morphological terminology and identify plants by sight and by using a dichotomous key. Along the way, we hope to alleviate plant awareness disparity (PAD), formerly known as plant blindness, by opening our students’ eyes to the plants surrounding us (Balas & Momsen, 2014; Jose et al., 2019; Krosnick et al., 2018; Parsley, 2020).

Plant taxonomy is often assessed through a plant collection project which authentically familiarizes students to the botanical terminology, dichotomous keys, pressed plants, and more recently iNaturalist (Krimmel et al., 2021; Liston & Struwe, 2018). Plant identification has traditionally been a very tactile experience where students manipulate specimens, experiencing the textures and orientations of leaves, stems, flowers, and fruit. There are 620 plant families made up of 16,167 genera, and approximately 391,000 species globally with new species being discovered on a continuous basis. While it is impossible to memorize them all, students can make use of the skills and knowledge they acquire through plant taxonomy courses and published dichotomous keys to identify nearly every species on the planet. Dichotomous keys are an incredible tool that taxonomists of all fields of biology and beyond can use to help narrow the specific taxonomic identity of an organism of interest. By providing relatively simplified pairs of choices that either succeed or fail to fit the description of the organism of interest, someone unfamiliar with a specific taxon may be led to the exact species of plant using only simple dissection tools. iNaturalist is designed to help anyone “Connect with Nature” by providing a one-stop shop for organismal identification, social media, locality data, and wildlife photography. It was designed with ease of use in mind, but also to aid scientists with data collection. With over 300,000 active users, iNaturalist is one of the most successful citizen science projects and has been used in over 120 peer-reviewed scientific papers (https://www.iNaturalist.org/stats). Anyone can upload photos, videos, or audio to iNaturalist through the app or website, and using automated image recognition, they can receive a tentative identification of an organism. While helpful, these suggested identifications can lack specificity and require experts or the use of a dichotomous key to properly identify to genus or species level (Heberling & Isaac, 2018; Jones, 2020). However, iNaturalist’s ability to identify organisms using its automated recognition software will only improve as more observations are made. Teaching identification using dichotomous keys and hands-on experiences will remain a core part of all plant taxonomy courses until the software is perfected. iNaturalist provides the perfect platform for university courses to explore and expand citizen science efforts.

iNaturalist does not use traditional tactile methods for identification. However, the ability to work with a global online community connects scientists, amateurs, and students in a way that manual keying cannot. Using a dichotomous key in an in-person lab setting can be collaborative as students work together on the same plant to use the provided key, but iNaturalist allows for a more diverse collegiality between different people, institutions, and regions. As the course at UGA included both Plant Biology major and non-major students, iNaturalist provided a way to encourage citizen science without overwhelming students with the advanced botanical vocabulary often required to use dichotomous keys in traditional classrooms.

Building upon past projects, we aimed to implement an improved plant collection project that achieved three main objectives for our students:

- Students will be able to correctly apply plant morphological terminology,
- Students will be able to identify plants to the species level using dichotomous keys, and
- Students will be able to create high-quality plant observations on the iNaturalist platform that are useable for future scientific research.

## Methods

iNaturalist has over 36.6 million observations recorded for plants alone. While many observations are unaffiliated with specific groups, there is a feature in iNaturalist where university professors can create projects to which their students can contribute. Data from eight different university plant collection projects, completed between March 2020 and May 2021, were downloaded from their project pages on iNaturalist (Table 1). Between the eight projects, there were a total of 6,911 observations. The raw data was filtered to exclude non-research grade observations (lacking 2 or more confirmed identifications to species level), non-plant observations (animals, fungi, etc.), and observations lacking locality data. In many of the projects, students were motivated to submit more than their required number of observations or resubmit observations originally deemed below par. While this encouragement may reduce the percentage of useful observations in a project, it is pedagogically sound to allow students revision and exploration in student-centered projects. Plant taxonomy university projects were identified from Rutgers University (RU), University of California Berkeley (UCB), Oregon State University (OSU), and the Connecting Students to Citizen Science and Curated Collections (CSCSCC) Project which includes classes from Arkansas State University, University of Michigan, Central Michigan University, and Middle Tennessee State University. These university projects were chosen based on their similar rubrics and larger number of observations. A stratified random sample of 160 observations was taken from each selected university’s project excluding our own. The UGA project had a total of 640 observations after filtering, all of which were included in the analyses for a grand total of 1280 observations across all universities. Observations were blinded by removing all identifying information except the iNaturalist observation ID#, URL to the observation, and species identification before being scored using our custom rubric (Table 2).

**Table 1.**
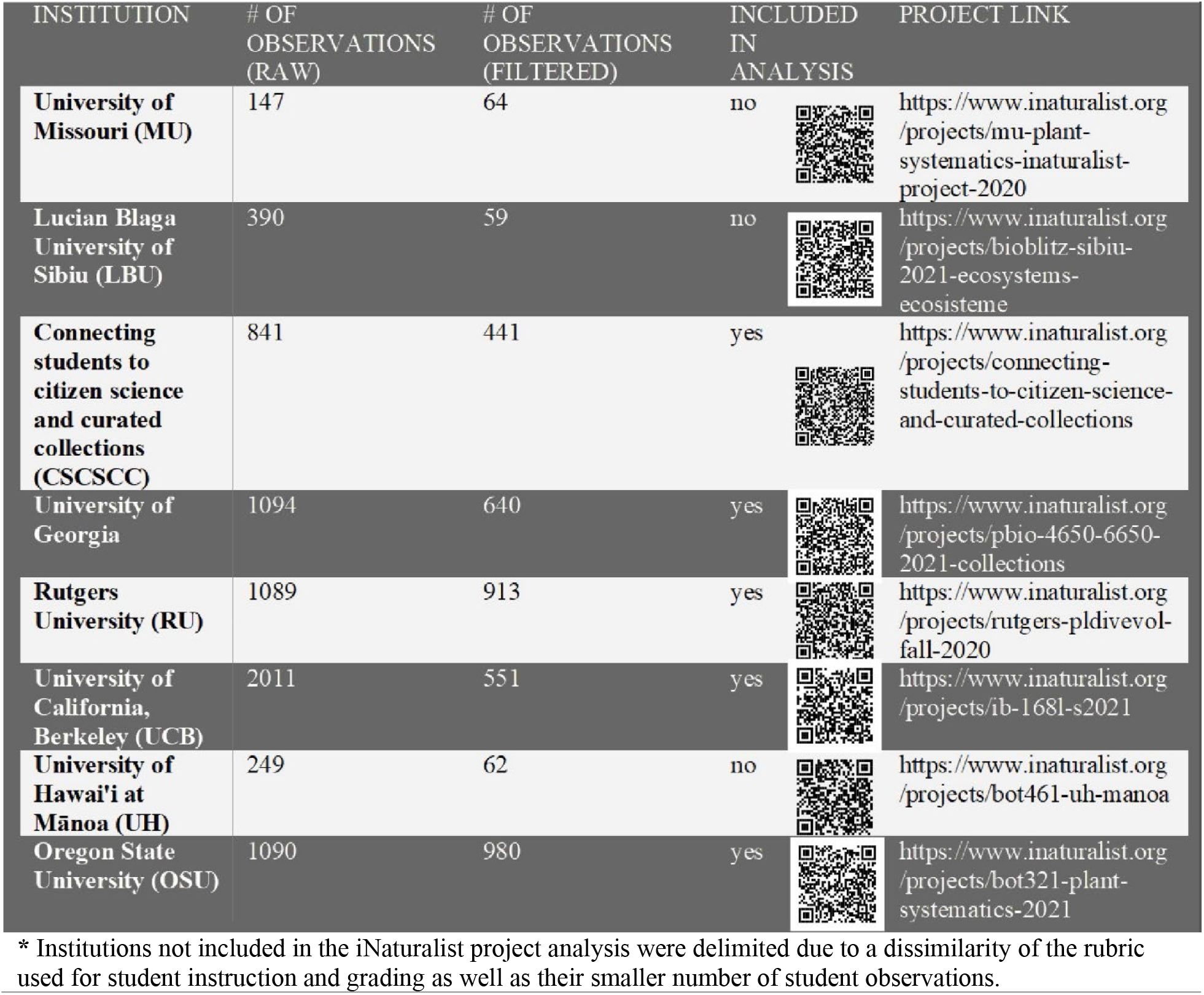
Data sources.

| INSTITUTION | # OF<br>OBSERVATIONS<br>(RAW) | # OF<br>OBSERVATIONS<br>(FILTERED) | INCLUDED<br>IN<br>ANALYSIS | PROJECT LINK |
| --- | --- | --- | --- | --- |
| **University of Missouri (MU)** | 147 | 64 | no | https://www.inaturalist.org/projects/mu-plant-systematics-inaturalist-project-2020 |
| **Lucian Blaga University of Sibiu (LBU)** | 390 | 59 | no | https://www.inaturalist.org/projects/bioblitz-sibiu-2021-ecosystems-ecosisteme |
| **Connecting students to citizen science and curated collections (CSCSCC)** | 841 | 441 | yes | https://www.inaturalist.org/projects/connecting-students-to-citizen-science-and-curated-collections |
| **University of Georgia** | 1094 | 640 | yes | https://www.inaturalist.org/projects/pbio-4650-6650-2021-collections |
| **Rutgers University (RU)** | 1089 | 913 | yes | https://www.inaturalist.org/projects/rutgers-pldivevol-fall-2020 |
| **University of California, Berkeley (UCB)** | 2011 | 551 | yes | https://www.inaturalist.org/projects/ib-1681-s2021 |
| **University of Hawai'i at Mānoa (UH)** | 249 | 62 | no | https://www.inaturalist.org/projects/bot461-uh-manoa |
| **Oregon State University (OSU)** | 1090 | 980 | yes | https://www.inaturalist.org/projects/bot321-plant-systematics-2021 |

\* Institutions not included in the iNaturalist project analysis were delimited due to a dissimilarity of the rubric used for student instruction and grading as well as their smaller number of student observations.

**Table 2.** Scoring rubric designed to reward observations with photographs with details relevant to keying plants to species

| POINT<br>VALUE | TRAIT |
| --- | --- |
| 1 | Photo(s) show leaf |
| 1 | Photo(s) show habitat (3 points of reference to provide context, e.g., surrounding vegetation, light levels, anthropogenic structures) |
| 1 | Photo(s) show growth habit (tree, shrub, vine, herb, etc.) |
| 1 | Photo(s) show leaf arrangement (opposite, alternate, whorled) |
| 1 | Photo(s) show mature reproductive structures (flower, fruit, cone, spores) |
| 1 | If Angiosperm: Photo(s) show enough characters to create at least 3/5 parts of a floral formula (flower shape[bilateral/radial], # sepals, # petals, # anthers, ovary superior/inferior); or fruit shows cross section<br>OR<br>If Gymnosperm: Photo(s) show male and female reproductive structures<br>OR<br>If Fern: Indusium clearly visible<br>OR<br>If Lycopod: Sporing structure clearly visible |
| 1 | Photo(s) contain ruler or coin for scale (hands and random objects do not count) |

|  |  |
| --- | --- |
| POINT VALUE | TRAIT |
| 1 | Photo(s) show leaf |
| 1 | Photo(s) show habitat (3 points of reference to provide context, e.g., surrounding vegetation, light levels, anthropogenic structures) |
| 1 | Photo(s) show growth habit (tree, shrub, vine, herb, etc.) |
| 1 | Photo(s) show leaf arrangement (opposite, alternate, whorled) |
| 1 | Photo(s) show mature reproductive structures (flower, fruit, cone, spores) |
| 1 | If Angiosperm: Photo(s) show enough characters to create at least 3/5 parts of a floral formula (flower shape[bilateral/radial], # sepals, # petals, # anthers, ovary superior/inferior); or fruit shows cross section<br>OR<br>If Gymnosperm: Photo(s) show male and female reproductive structures<br>OR<br>If Fern: Indusium clearly visible<br>OR<br>If Lycopod: Sporing structure clearly visible |
| 1 | Photo(s) contain ruler or coin for scale (hands and random objects do not count) |

In designing the rubric, the Botany Depot website was searched for extant rubrics. Aaron Liston published a rubric; soon after, professors from the CSCSCC published a similar rubric on their own website. All projects included in the analyses were based upon either the Liston or CSCSCC rubrics which share similar grading criteria and learning objectives (Krimmel et al., 2021; Liston & Struwe, 2018). Students at UGA were shown how to make observations in an iNaturalist project and given a rubric at the beginning of the semester (Supplementary Document 1). The rubrics made available from all colleges’ projects were based on the same premise of wanting to create research-grade photograph collections. In addition to the requirements of the original rubrics, the UGA rubric required student observations to include a minimum of four photographs of the plant with at least one image containing a coin or ruler for scale. Images were required to show leaf shape, leaf arrangement, stem, reproductive structures (fruit or flowers), and overall habit of the plant (how it grows). Every observation was required to contain all the necessary information to make a proper herbarium label in the observation notes section. Undergraduate students were required to submit 20 observations and graduate and honors students were required to submit 30 observations. Lastly, each student was required to key out (identify using a dichotomous key) two of their classmates’ observations and three of their own and post the steps as a comment on iNaturalist.

Scoring was completed using a robust rubric designed to assess the ability of an observation to be identified using a dichotomous key and utilized by researchers (Table 2). The rubric was designed to reward observations that include the level of detail often required for identifying a plant to the species level using a dichotomous key. Orientation, count, and tiny details such as presence or size of glands, hairs, or vein patterns are not infrequently the deciding factor in a dichotomous key. And while many plants like lilies and roses may have large, easily observable flowers, there are far more flowering plants passed by daily with much smaller flowers (e.g., grasses). The more of these fine details that were captured in the observation photos, the higher the score and the more utility that observation has for future research. Observations that were pressed, cultivated, completely plucked from where they were growing with no indication of where they came from, or containing all blurry photos were given a score of zero. Once all observations were scored, an ANOVA was performed in R to compare scores across universities.

## Results

Each of the five universities or consortiums sampled included observations that represented the full range of the rubric, from zero to seven points, and all average scores were above 3.5 points. Despite UGA having more observations than the other groups included in analysis, the standard deviation of UGA’s observations (1.51) was in the middle of the other four universities (OSU: 1.31, UCB: 1.47, CSCSCC: 1.75, RU: 1.80). OSU had the highest mean score (5.26) followed by UGA (5.07). The other three projects’ scores were significantly (p < 0.05) lower with UCB scoring an average of 4.27, CSCSCC scoring an average of 3.96, and RU scoring an average of 3.76 (see Figure 1). The median and mode of UGA and OSU were 5 points and 6 points respectively. CSCSCC and UCB showed median and mode of 4 points and 5 points respectively, and RU showed 4 points for both measures.

**Figure 1.**
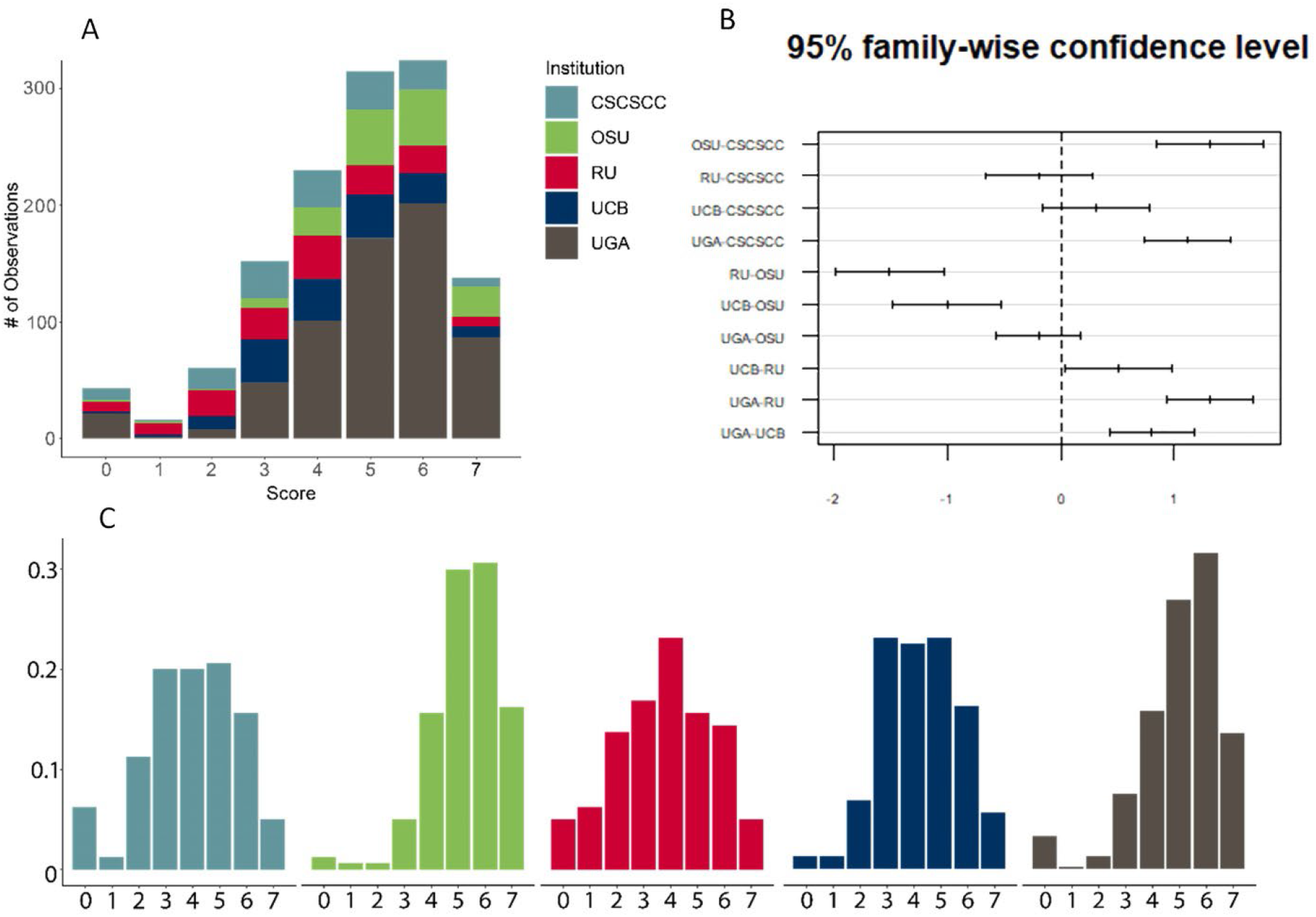
**A)** Total number of observations for each score category, colors indicate institution. **B)** The average change in mean response (mean score) when comparing between institutions. **C)** Percent of observations for each score, color indicates institution.

During scoring, most of the points missed resulted from a lack of photographic detail of reproductive structures, overall habitat, and failing to include a standardizable scale in at least one photograph. The ANOVA revealed that the observations in our project (UGA) had significantly higher scores than CSCSCC, UCB, RU, and all institutions included combined (p<0.001). However, our observations were not significantly better than the ones from OSU (p=0.64, Figure 1). OSU also had significantly higher scores than all other institutions except UGA (see Figure 1).

## Discussion

The global pandemic forced lab- and lecture-based classes to move online in the spring of 2020. Plant taxonomy has rarely, if ever been offered in an online format. At UGA this required extensive updating and modification over a two-week timespan in March 2020 and throughout the spring 2021 semester; plant taxonomy is not offered in fall semesters. One of the greatest losses of moving to remote teaching was the inability to discuss with students about discoveries they made for plants that they have in-hand. Using iNaturalist, students can go for a walk around their neighborhood, take pictures of any plant they happen across and receive a best guess on the identification immediately. This jump-starts learning about the plant they came across and may choose to include in their collection project. After adding a plant observation to the iNaturalist app, students could easily add it to the class project for instructors to review, comment on, confirm, or reject.

Through the semesters, we found several negative effects on our students’ learning as a result of remote instruction and reliance on iNaturalist over physical keying. Dichotomous keys require practice and experience to utilize properly. If students are introduced to iNaturalist in a classroom setting before becoming comfortable with identifying specimens using dichotomous keys, the automatic identification capabilities can be used to circumvent exercising keying skills. Students may no longer feel the need to learn the process of keying out specimens to get a potential identification if there is an app giving them a putative answer. The problem remains that app-based identification is far from perfect. iNaturalist was only able to identify plants to the correct species 35% of the time and provided entirely incorrect, misleading IDs 16% of the time (Jones, 2020). Even if the ID is only to family level, if it is correct then students need not utilize the key to family in the flora being used for identification (i.e., if the app’s ID is partially correct, it is detrimental to student learning).

However, different aspects of student learning were improved through use of iNaturalist. Making plant taxonomy a student-centered learning experience rather than content-centered course should be a priority. Having clear, learner-oriented objectives, instructions, and rubrics available to students at all times gives the students the power to succeed while minimizing stress (Kuiper et al., 2015). Most plant taxonomy courses occur in the spring to maximize the success of observing reproductive structures; in North America, this means that the semester will begin in January and end in late April or May. Depending on how and when the project is introduced, students may have three weeks to two months of prime collecting season. Factoring in the available time for observations, instructors may consider reducing requirements such as the number of observations or pressed specimens, but further research is required to support this assertion. The instructor created a revised rubric based on student feedback to use in future semesters (Supplementary Document 2). Even with only 20 observations required for undergraduates and 30 observations required for graduate and honors students, the instructors were approached multiple times with concerns of the collection project being overwhelming. Providing students with extra opportunities to make observations for the project during optional field trips scheduled outside of normal class time seemed to alleviate most of these issues.

One of the primary goals of plant taxonomy courses is to give students the skills to take nearly any plant they encounter through a dichotomous key and identify it to the species level. Distinguishing a specimen using a key often requires dissecting reproductive structures (flowers and fruits) at a fine level with great attention to detail. iNaturalist provides a less-invasive option for recording the presence of rare and endangered species. Unlike pressing plants, iNaturalist observations are non-destructive and can provide a plethora of habitat, ecological, environmental, and climate information. However, iNaturalist is not a replacement for physical collections. Herbaria and the physical plant collections they contain are invaluable as a source of morphometric and genetic data (Bieker & Martin, 2018; Fritsch et al., 2018; Miller-Rushing, Primack, Primack, & Mukunda, 2006; Weaver, Ng, & Laport, 2020). However, photographs of living specimens do provide additional insights especially when exact locality, date, and time information is included, and photos showcase three dimensional aspects not captured in the pressing process. Some herbaria have even begun integrating photography into their collections to facilitate this new source of information (Gómez-Bellver et al., 2019; Heberling et al., 2019; Heberling & Isaac, 2018).

## Conclusion

The changes made to the UGA plant collection project and rubric significantly improved the quality of iNaturalist observations with regards to students’ ability to be run through a dichotomous key to species level identification compared to all other universities combined. The rubric required student observations to include more and higher quality details for each observation. These observations are more likely to be suitable for use in research studies (Barve et al., 2020; Echeverria et al., 2021; Gazdic & Groom, 2019; Horn et al., 2018; Schiller et al., 2021). It is our hope that moving forward plant taxonomy courses make use of the rubric and continue improving upon it in a student-centered manner.

## Supplementary Document 1

### iNaturalist Plant Collection Guidelines

Collecting, identifying, and preparing plants for herbaria records are important skills to learn as a plant taxonomist. You will be expected to complete a digital plant collection for a portion of your course grade. All students will complete a collection with 20 plants (Undergraduates) or 30 plants (Honors/Graduate Students) worth 200 or 300 points, respectively.

### Objectives

- Learn how to identify plants using a dichotomous key
- Develop the ability to make informative collection notes and records
- Gain awareness for the plants around you
- Learn how to integrate photography into herbarium collections

### Due Dates

- **April 14**^**th**^ **5pm** - 10 observations due to iNaturalist class project (15 observations for honors/graduate)
- **May 5**^**th**^ **5pm** – 10 additional observations due to iNaturalist class project (15 additional observations for honors/graduate); **All labels and ID comments (iNaturalist) due**

### Plant Collection Requirements

- Collect the specimen with a series of photos (4 minimum)
  - You must include photos clearly depicting the flower, leaf, arrangement, stem, and inflorescence, as well as fruit, if applicable. Be sure to document any range in features.
  - Include a quarter or scale bar in at least one photo per observation
  - Angiosperms suggested, but not required

- Specimens **<u>must</u>** include reproductive structures (flowers, fruits, or sporangia) along with the vegetative portion of the plant.
  - If photos were taken before March 24^th^ reproductive structures are not required. However, enough phenotypic characters for exact species identification are required.

- Only native or widely introduced (naturalized) plants should be collected. **Do not collect cultivated plants**.
- All specimens should be uploaded to the class iNaturalist Project. Further instructions for this are in the “Using iNaturalist” document.
- Virtual labels should be prepared and included with your submissions to iNaturalist. See below for more information on label requirements.
- **You must comment on a minimum of 10 (Undergraduate) or 15 (Honors/Graduate) observations (not your own) within the class iNaturalist project** and provide a confirmation of the observer’s ID
- *Only 1 of your ID confirmation/suggestions may be on an observation made by the planttax1 or mzenoble accounts*

### Collecting Plants in the Field

- Choose plants that are well developed and not diseased.
- The entire plant should be photographed; never record just one flower or leaf (unless that is the entire plant)!
- Avoid collecting specimens lacking flowers, fruits, or other reproductive structures; such material will be difficult to identify.
- If not making observations in the field be sure to record ALL relevant information outlined in the Virtual Label Requirements section

### Virtual Label Requirements aka iNaturalist Notes Section

The label should include all the following information about each specimen:

1. Family
2. Genus
3. Species
4. Country*
5. State*
6. County*
7. Nearest major road/city
8. GPS Coordinates*
9. Elevation*
10. Date photographed*
11. Habitat type
12. Associated species
13. Additional characters (see bullet point below)
14. Collector ID (your myID)
15. Specimen ID number
16. Any additional notes

**(*) Fields with asterisks indicate fields that should be automatically recorded if photos are taken within the iNaturalist app. If you take photos not through the app this information may not be recorded automatically**. If you need assistance with this, ask your instructors for help.

- iNaturalist was not designed with integration with herbarium labels in mind, so much of the label information required for proper herbarium specimens will not have its own field for entry. **Include all the label information in the “Notes” section of the observation submission form unless submitting via computer**.
- Good (and bad) examples may be found by browsing the herbarium records through SERNEC.
- **Additional characters** - Information concerning the plant that won’t be evident in the photos, such as scent, bark characteristics, local abundance, pollinators, etc.

### Plant identification

- Plants should be keyed/identified using the Weakley 2020 Flora
- You are welcome to seek assistance from each other or from your instructors
- Additionally, if you are completely stumped as to a specimen’s identification you can attempt to use the Seek app to get a preliminary ID. You must still fully key out the 3 specimens you collect regardless of whether you use the Seek app.

**NOTE:** Seek is typically great for familial level identification, but not ideal for species level identification.

### Collection submission (Word document to ELC by 5/5/2021 @5pm)

- Include your iNaturalist username, your actual name, and your lab section (Page 1)
- Include a list of the plants in the collection, organized alphabetically by family, then by genus, then by species (Page 1)
- List the 10/15 observations you commented on within the iNaturalist project and the username(s) of the person that collected them
- Write down the steps taken to confirm identification for 3 specimens **you** observed using the dichotomous keys in Weakley’s Floras (Page 2+)

### Collection Grading Sheet

- You must include a photos series of your specimen (include a photo of the flower, leaf, stem, and inflorescence or fruit, if applicable)
- 20 species of vascular plants (for Undergraduates)
- 30 species of vascular plants (for Honors/Graduates)
- Angiosperms suggested, but not required
- Minimum of 10 families represented in your observations, 15 for graduate/honors **OR** do a case study of a single family (all specimens from the same family; i.e. Fabaceae)
- Only native or naturalized plants should be collected. **Do not collect cultivated plants**.
- Specimens **<u>must</u>** include fertile material (flowers, fruits, cones, or sporangia) along with the vegetative portion of the plant
- All label information must be included in the notes section or custom fields of every submission to iNaturalist
- You must comment on 10 (Undergraduate) or 15 (Honors/Graduate) of your classmates’ observations and provide confirmation or alternative IDs; **2 of these comments (3 for graduate/honors) must include the steps in the dichotomous keys** you used to reach your conclusion (do not copy and paste these from the observer)

### Undergraduate Grading Rubric

Each specimen will be graded according to the following criteria:

1. Adequate photos – properly focused, not grainy or blurry, includes coin for scale (0.5 per photo, 2/specimen, 40 total)
2. ID of species – (2 each, 40 total)
3. Correct ID of genus – (1.5 each, 30 total)
4. Correct ID of family – (1 each, 20 total)
5. Label information complete – (2 each, 40 total)
6. ID Comments on classmate observations – (1 each, 10 total)
7. Keys to 3 of your specimens – (4 each, 12 total)
8. Key to 2 of your classmates’ observations – (4 each, 8 total)

**Total 200 points**

### Honors/Graduate Grading Rubric

Each specimen will be graded according to the following criteria:

1. Adequate photos – properly focused, not grainy or blurry, includes coin for scale (0.5 per photo, 2/specimen, 60 total)
2. ID of species – (2 each, 60 total)
3. Correct ID of genus – (1.5 each, 45 total)
4. Correct ID of family – (1 each, 30 total)
5. Label information complete – (2 each, 60 total)
6. ID Comments on classmate observations – (1 each, 15 total)
7. Keys to 5 of your specimens – (4 each, 20 total)
8. Key to 3 of your classmates’ observations – (3.3 each, 10 total)

**Total 300 points**

## Supplementary Document 2

### Assignment: Plant Collection Project using iNaturalist and Pressed Plants

#### Due

- **April 5**^**th**^ **5pm** - 10 observations due to iNaturalist class project (15 observations for honors/graduate); 10 pressed plants (15 for honors/graduate) due
- **May 3**^**rd**^ **5pm** – 10 additional observations due to iNaturalist class project (15 additional observations for honors/graduate); **All labels (iNaturalist and physical) and ID comments (iNaturalist) due;** return plant presses

### Submission Instructions

Submit a .xlsx or .csv file containing the 20 (undergraduate) or 30 (honors/graduate) observations you made on iNaturalist in the class project (template on eLC). Bring a folder containing all your specimens to lab by the April 5th due date. Submit all physical specimens including properly formatted labels for grading by the final due date. Follow the Using iNaturalist document for further instructions.

### Purpose

Students will be able to identify plants to the species level using a dichotomous key and gather relevant details required for herbarium grade plant observations which reinforces the knowledge and critical thinking skills required for proper plant identification. This summative assessment determines if you can correctly apply the skills and knowledge gained throughout the course to learning outcomes 2 and 3 (on the syllabus). This is a chance to explore nature and notice plants, flowers, and fruits that you likely often walk past but never noticed and to learn if they are useful for people through a collaborative and social media-like platform (iNaturalist).

### Skills

The purpose of this assignment is to help you practice the following essential skills:

Photograph plants during their reproductive stage

Record notes that aid in the identification of the plant

Press plants using quality techniques

Use the Weakley Flora to identify the plant to the species level

### Knowledge

This assignment will also help you become adept at applying the following important concepts and pedagogical knowledge to real-world tasks:

Apply deductive reasoning and critical thinking to problem solving by combining prior knowledge and physical, observable traits into a proper, tentative identification.

### Task

Press, dry, and prepare for mounting 10 plants (15 for honors/graduate) collected throughout the semester. Create properly formatted herbarium specimen labels and submit all your specimens by the assigned due date.

Properly formatted herbarium labels include the following:

- *Family*
- *Genus*
- *Species*
- *Country*
- *State*
- *County*
- *Nearest major road/intersection*
- *GPS coordinates*
- *Elevation*
- *Data collected*
- *Habitat type*
- *Associated species*
- *Additional characters*
- *Collector ID#*
- *Specimen ID#*

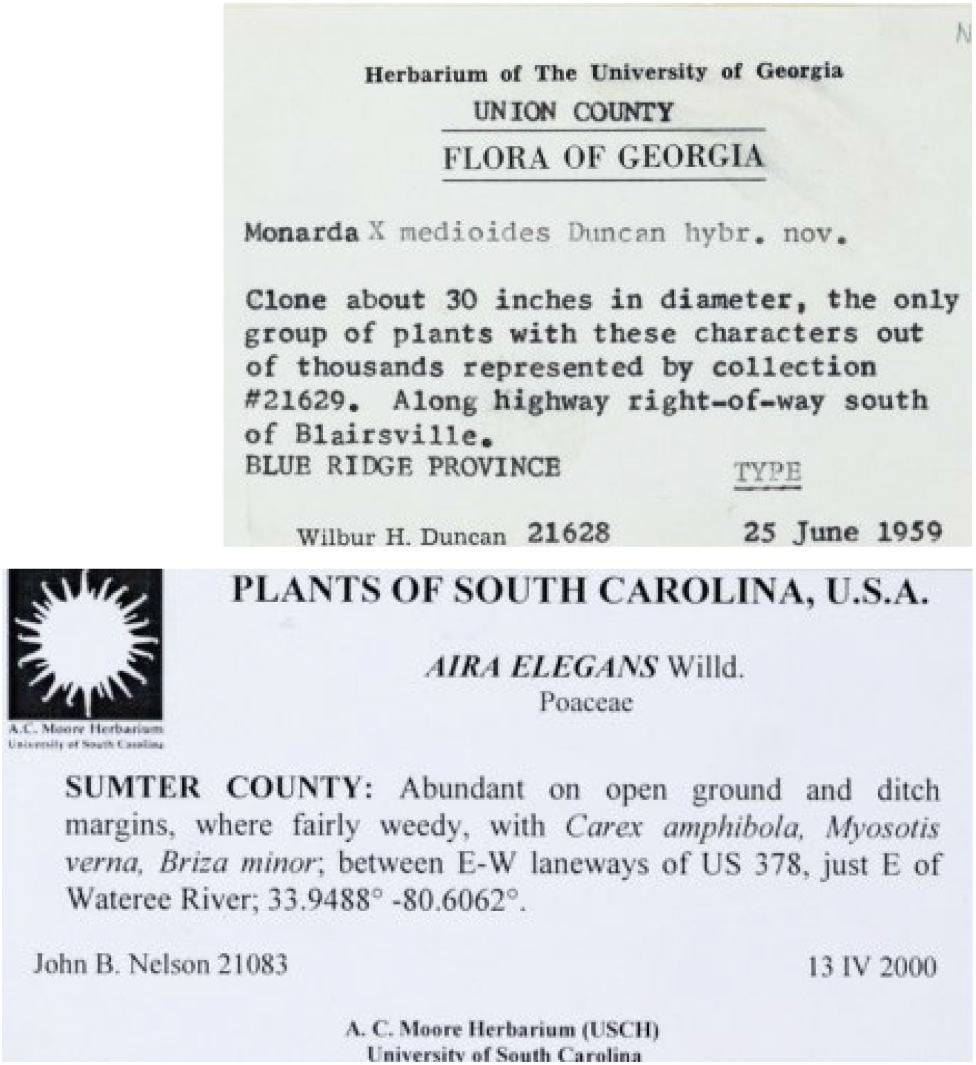

Take multiple photos (minimum of 4) of the plant that include a standard coin or ruler for scale in as many photos as possible.

### Photos should include the following

- *Closeups of the flower parts* [if you can take pictures from the top, side, and bottom of the flower this usually gets everything]
- *A single, whole leaf* [be careful to be sure you can see the node, so you aren’t taking a portion of a compound leaf]
- *Leaf arrangement* [opposite, alternate, whorled]
- *The fruit* [including a horizontal cross section if possible]
- *The entire plant* [to show growth habit: vine, shrub, tree, herb, etc.]
- *The habitat where the plant is growing* [the plant may be too small to be clearly visible in this photo]

Based on your images and the plant, add a list of appropriate terminology learned throughout the course to those to the notes section of your observation on iNaturalist.

Use the Weakley Flora to key out the specimen first to family and then species.

List the steps you took within the key in the comments section of your observation, including a brief summary of the trait that you used to decide on each step. All the above will be practiced during lab.

### Criteria for Success

A successful project will meet all the requirements as outlined below, during the first lab, collecting field trips, and the rubric posted on eLC.

If you need help on this project, the following options are available to you:

- You may ask any of the instructors to look at your observations prior to the due dates above, during lab field trips, or during office hours.
- You can confirm or reject your classmates’ observations on the class iNaturalist page by keying their observations out using the Weakley Flora at any time during the semester for additional practice (instructors will be happy to confirm your IDs).
- You may look at the 2021 PBIO 4650/6650 iNaturalist project which contains multiple sample observations made from the prior semester that should be properly identified, include field notes, and keying steps. Note: The observations in this project were made with a slightly different rubric and may be missing one or more of the required components from this assignment prompt.

We strongly encourage you to get creative and travel anywhere you want during the semester to explore and make observations of unusual plants in unique locations/habitats [Do not trespass, physically remove any plant material from property you do not have permission to collect on, or make observations of cultivated plants for this project].

Physical collecting outside of guided field trips (by the instructors) is allowed for non-threatened and/or endangered plants if you have proof of written landowner permission. This written permission must be submitted alongside the pressed specimens.

Extra Credit will be given for each unique species identified across the entire class iNaturalist project (e.g., you are the only person to observe the species Cypripedium acaule (Pink Lady’s Slipper orchid) in the entire class). Full point details in the grading rubric below. This does not apply to physically pressed specimens.

### Requirements

1. You must make observations of a total of 20 species of vascular plants (for Undergraduates) or 30 species of vascular plants (for Honors/Graduates). 10 of the iNaturalist observations (15 for honors/graduate) must be linked to physical specimens you collect, press, dry, and mount. We will teach you how to do the physical collections in lab and provide opportunities to collect during field trips.
2. All observations must be of wild, naturally occurring plants. No cultivated plants! Weeds in planted areas are okay. The native plant garden in the State Botanical Gardens of Georgia is considered cultivated. If an observation’s GPS coordinates put it overlapping with that portion of the garden it will not count for this assignment.
3. You must comment on a minimum of 10 (Undergraduate) or 15 (Honors/Graduate) observations (not your own) within the class iNaturalist project and provide a confirmation of the observer’s ID [please do not just press “I agree” without keying it out first unless you are 100% certain]
4. Only 1 of your ID confirmation/suggestions may be on an observation made by the instructor accounts
5. 5.Record details about each plant as explained in the Observation Notes section below
6. Upload photos (minimum of 4) of the plant to the class iNaturalist project that include a standard coin or ruler for scale in as many photos as possible. Cropped versions of the same photo only count as one photo. Photos from the internet is considered academic dishonesty. Photos should include the following:
  - Closeups of the flower parts [if you can take pictures from the top, side, and bottom of the flower this usually gets everything]
  - A single, whole leaf [be careful to be sure you can see the node, so you aren’t taking a portion of a compound leaf]
  - Leaf arrangement [opposite, alternate, whorled]
  - The fruit [including a horizontal cross section if possible]
  - The entire plant [to show growth habit: vine, shrub, tree, herb, etc.]
  - The habitat where the plant is growing [the plant may be too small to be clearly visible in this photo; scale not required]
7. Use the Weakley Flora to key out each specimen first to family and then species.
8. List the steps you took within the key in the comments section of your observation, including a brief summary of the trait that you used to decide on each step. All the above will be practiced during lab field trips.

### Observation Notes

iNaturalist records a lot of information automatically if you allow it to that is relevant to herbarium specimens but it doesn’t get everything.

- iNaturalist should automatically record the country, state, county, GPS coordinates, elevation, date photographed, and your collector ID if you make observations on your phone. [Note: Unfortunately, if you make observations on your computer, you will likely have to manually enter this information]
- You must manually record the nearest major road/intersection, the habitat type, any associated species, additional characteristics, collector ID, and the specimen ID number.
- Based on your images and the plant, add a list of appropriate terminology learned throughout the course to those to the additional characters section of each observation
- Additional characters may include but are not limited to information concerning the plant that won’t be evident in the photos, such as scent, bark characteristics, local abundance, pollinators, etc.

### Scoring Rubric

#### Pressed specimens 50 (5pt each)

- *Pressed to GA herbarium standards* [see How to Make Herbarium Specimens document on eLC] *(2pt)*
- *Specimen label (3pt)*
  - *Proper formatting (1pt)*
  - *Contains all relevant information (1pt)*
  - *iNaturalist observation # (1pt)*

### Photos 40 (2pt each)

- *Reproductive structures (0*.*4pt total)*
  - *components to complete 4/5 of a floral formula [if in flower, 0*.*1pt each] OR*
  - *entire fruit and fruit placentation [if in fruit, 0*.*2pt each]*

- *Vegetative structures (0*.*4pt total)*
  - *Entire leaf (0*.*2pt)*
  - *Leaf Arrangement (0*.*2pt)*

- *Growth Habit (0*.*4pt)*
- *Habitat (0*.*4pt)*
- *Scale present (0*.*4pt maximum)*
  - *In all photos [except habitat] (0*.*4pt)*
  - *In some photos (0*.*2pt)*
  - *In 1 photo (0*.*1pt)*
  - *In no photos (0pt)*

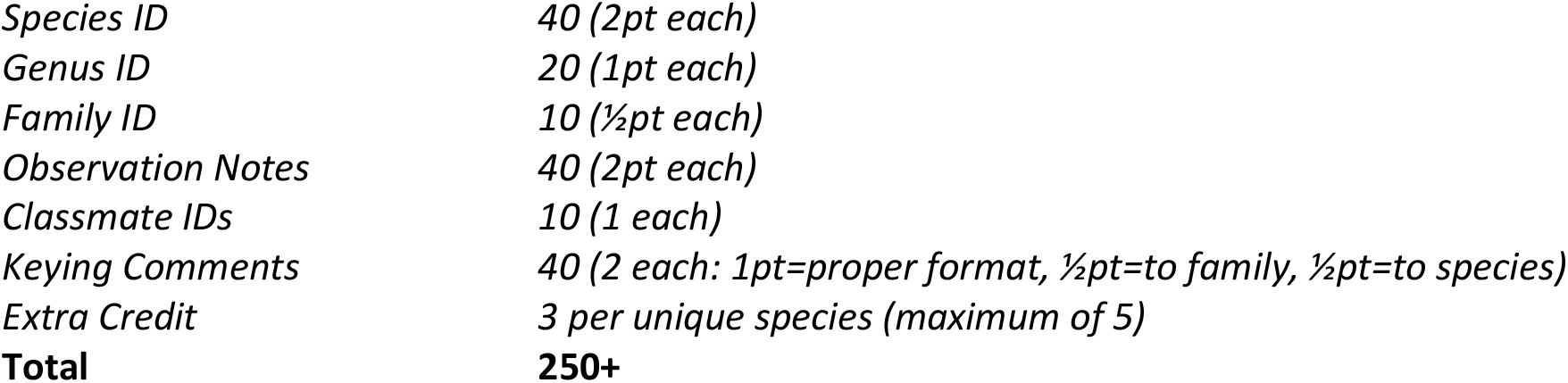

## References

Balas, B., & Momsen, J. L. (2014). Attention “blinks” differently for plants and animals. CBE Life Sciences Education, 13(3), 437–443. https://doi.org/10.1187/cbe.14-05-0080

Barve, V. V., Brenskelle, L., Li, D., Stucky, B. J., Barve, N. V., Hantak, M. M., McLean, B. S., Paluh, D. J., Oswald, J. A., Belitz, M. W., Folk, R. A., & Guralnick, R. P. (2020). Methods for broad-scale plant phenology assessments using citizen scientists’ photographs. Applications in Plant Sciences, 8(1), e11315. https://doi.org/10.1002/APS3.11315

Bieker, V. C., & Martin, M. D. (2018). Implications and future prospects for evolutionary analyses of DNA in historical herbarium collections. Botany Letters, 165(3–4), 409–418. https://doi.org/10.1080/23818107.2018.1458651

Echeverria, A., Ariz, I., Moreno, J., Peralta, J., & Gonzalez, E. M. (2021). Learning Plant Biodiversity in Nature: The Use of the Citizen–Science Platform iNaturalist as a Collaborative Tool in Secondary Education. Sustainability 2021, Vol. 13, Page 735, 13(2), 735. https://doi.org/10.3390/SU13020735

Fritsch, P. W., Nowell, C. F., Leatherman, L. S. T., Gong, W., Cruz, B. C., Burge, D. O., & Delgado-Salinas, A. (2018). Leaf adaptations and species boundaries in North American Cercis: implications for the evolution of dry floras. American Journal of Botany, 105(9), 1577– 1594. https://doi.org/10.1002/ajb2.1155

Gazdic, M., & Groom, Q. (2019). iNaturalist is an Unexploited Source of Plant-Insect Interaction Data. Biodiversity Information Science and Standards, 41. https://go.gale.com/ps/i.do?p=AONE&sw=w&issn=25350897&v=2.1&it=r&id=GALE%7CA646476433&sid=googleScholar&linkaccess=fulltext

Gómez-Bellver, C., Ibáñez, N., López-Pujol, J., Nualart, N., & Susanna, A. (2019). How photographs can be a complement of herbarium vouchers: A proposal of standardization. Taxon, 68(6), 1321–1326. https://doi.org/10.1002/tax.12162

Heberling, J. M., & Isaac, B. L. (2018). iNaturalist as a tool to expand the research value of museum specimens. Applications in Plant Sciences, 6(11), e1193. https://doi.org/10.1002/aps3.1193

Heberling, J. M., Prather, L. A., & Tonsor, S. J. (2019). The Changing Uses of Herbarium Data in an Era of Global Change: An Overview Using Automated Content Analysis. BioScience, 69(10), 812–822. https://doi.org/10.1093/biosci/biz094

Horn, G. Van, Aodha, O. Mac, Song, Y., Cui, Y., Sun, C., Shepard, A., Adam, H., Perona, P., Belongie, S., Google, C. 2, & Tech, C. (2018). The INaturalist Species Classification and Detection Dataset (pp. 8769–8778). http://www.inaturalist.org

Jones, H. G. (2020). What plant is that? Tests of automated image recognition apps for plant identification on plants from the British flora. In AoB PLANTS (Vol. 12, Issue 6). Oxford University Press. https://doi.org/10.1093/aobpla/plaa052

Jose, S. B., Wu, C. H., & Kamoun, S. (2019). Overcoming plant blindness in science, education, and society. Plants People Planet, 1(3), 169– 172. https://doi.org/10.1002/ppp3.51

Krimmel, E. R., Linton, D. L., Marsico, T. D., Monfils, A. K., Morris, A. B., & Ruhfel, B. R. (2021). Connecting Students to Citizen Science and Curated Collections. https://collectionseducation.org/

Krosnick, S. E., Baker, J. C., & Moore, K. R. (2018). The Pet Plant Project: Treating Plant Blindness by Making Plants Personal. American Biology Teacher, 80(5), 339–345. https://doi.org/10.1525/abt.2018.80.5.339

Kuiper, S. R., Carver, R. H., Posner, M. A., & Everson, M. G. (2015). Four Perspectives on Flipping the Statistics Classroom: Changing Pedagogy to Enhance Student-Centered Learning. Https://Doi.Org/10.1080/10511970.2015.1045573, 25(8), 655–682. https://doi.org/10.1080/10511970.2015.1045573

Liston, A., & Struwe, L. (2018). Botany Depot Activity: iNaturalist Observation Project. Botany Depot. https://botanydepot.com/2020/03/21/activity-inaturalist-observation-project-by-aaron-liston/

Miller-Rushing, A. J., Primack, R. B., Primack, D., & Mukunda, S. (2006). Photographs and herbarium specimens as tools to document phenological changes in response to global warming. American Journal of Botany, 93(11), 1667–1674. https://doi.org/10.3732/ajb.93.11.1667

Parsley, K. M. (2020). Plant awareness disparity: A case for renaming plant blindness. PLANTS, PEOPLE, PLANET, 2(6), 598–601. https://doi.org/10.1002/ppp3.10153

Schiller, C., Schmidtlein, S., Boonman, C., Moreno-Martínez, A., & Kattenborn, T. (2021). Deep learning and citizen science enable automated plant trait predictions from photographs. Scientific Reports 2021 11:1, 11(1), 1–12. https://doi.org/10.1038/s41598-021-95616-0

Weaver, W. N., Ng, J., & Laport, R. G. (2020). LeafMachine: Using machine learning to automate leaf trait extraction from digitized herbarium specimens. Applications in Plant Sciences, 8(6), e11367. https://doi.org/10.1002/aps3.11367

