## Supplementary Document 2 for "Emphasis on quality in iNaturalist plant collections enhances learning and research utility"

Apply deductive reasoning and critical thinking to problem solving by combining prior knowledge and physical, observable traits into a proper, tentative identification.

**Task:**

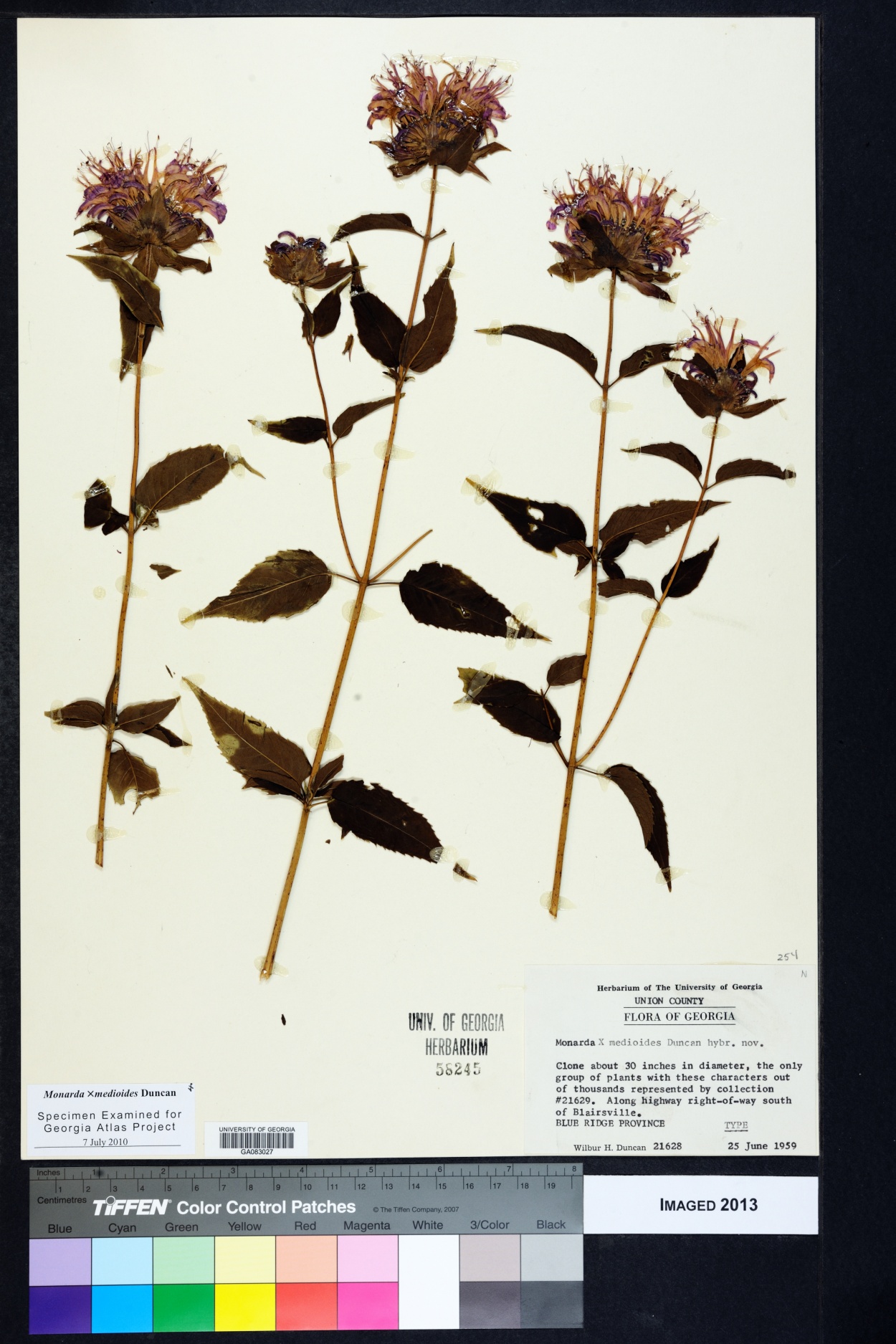
Press, dry, and prepare for mounting 10 plants (15 for honors/graduate) collected throughout the semester. Create properly formatted herbarium specimen labels and submit all your specimens by the assigned due date.

Properly formatted herbarium labels include the following:

- *Family*
- *Genus*
- *Species*
- *Country*
- *State*
- *County*
- *Nearest major road/intersection*
-
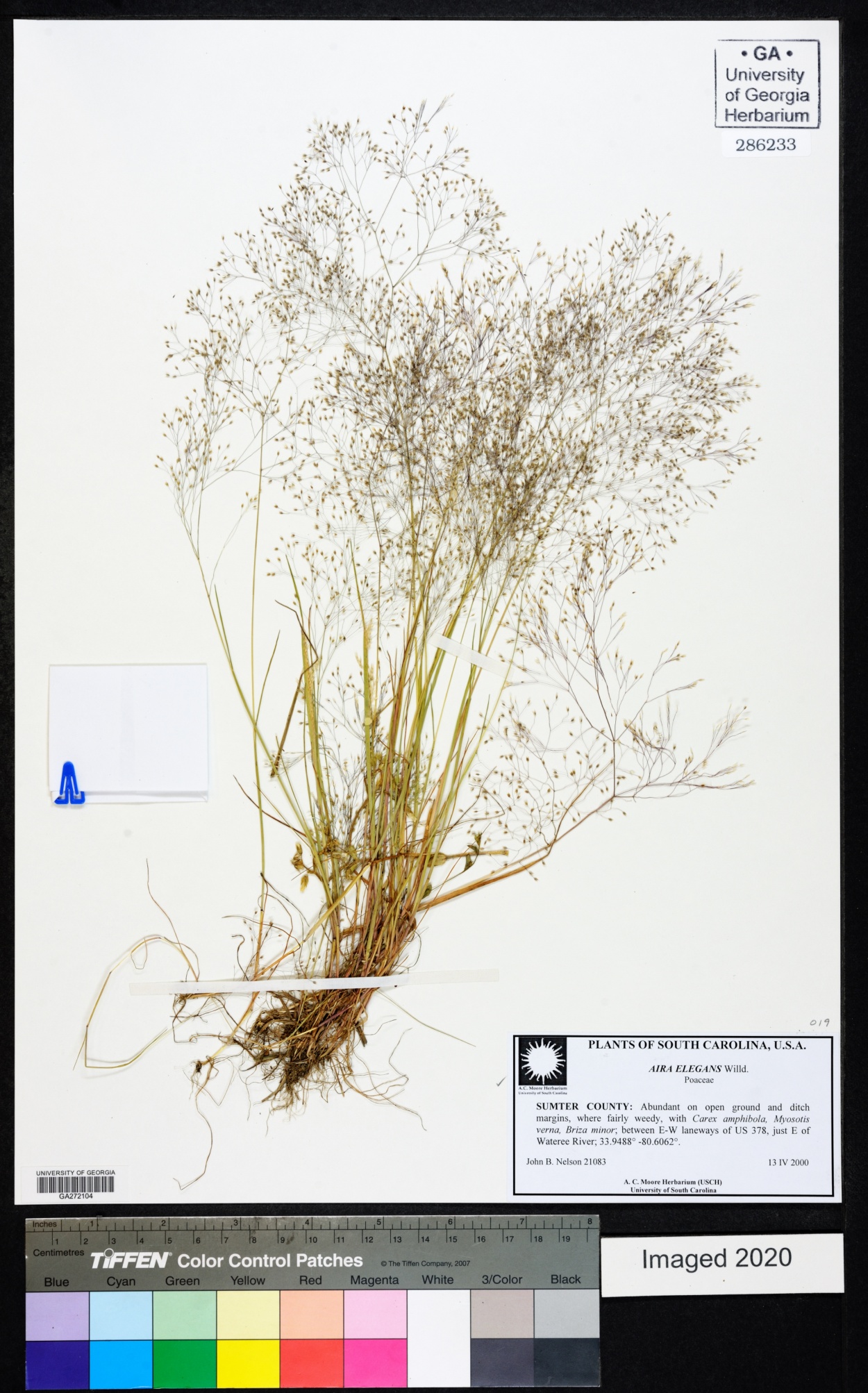
*GPS coordinates*
- *Elevation*
- *Data collected*
- *Habitat type*
- *Associated species*
- *Additional characters*
- *Collector ID#*
- *Specimen ID#*

*Photos	40 (2pt each)*

- *Reproductive structures (0.4pt total)*
  - *components to complete 4/5 of a floral formula [if in flower, 0.1pt each]
    OR*
  - *entire fruit and fruit placentation [if in fruit, 0.2pt each]*
- *Vegetative structures (0.4pt total)*
  - *Entire leaf (0.2pt)*
  - *Leaf Arrangement (0.2pt)*
- *Growth Habit (0.4pt)*
- *Habitat (0.4pt)*
- *Scale present (0.4pt maximum)*
  - *In all photos [except habitat] (0.4pt)*
  - *In some photos (0.2pt)*
  - *In 1 photo (0.1pt)*
  - *In no photos (0pt)*

*Species ID	40 (2pt each)*

*Genus ID	20 (1pt each)*

*Family ID	10 (½pt each)*

*Observation Notes	40 (2pt each)*

*Classmate IDs	10 (1 each)*

*Keying Comments	40 (2 each: 1pt=proper format, ½pt=to family, ½pt=to species)*

*Extra Credit	3 per unique species (maximum of 5)*

**Total	250+**
